# Assessment of *Hyptis suaveolens* Extract-Infused Candle Wax as a Mosquito Biorepellent

**DOI:** 10.64898/2026.07.22.740051

**Authors:** Enos Quarshie Tuleasi, Michaella Nyame Hayford, Beatrice Assantewaa

## Abstract

Neurotoxic chemicals, including pyrethroids, neonicotinoids, and chlorinated hydrocarbons, have been effective for mosquito control; however, their continued use has resulted in widespread insecticide resistance. This resistance has necessitated increased insecticide application, a phenomenon known as the pesticide treadmill, which poses significant risks to both environmental and human health. Consequently, there is a growing need for sustainable alternatives with minimal adverse impacts. Although plant-based repellents have shown promise, many are formulated as topical products that may be costly, inconvenient, or culturally unacceptable in some communities. Despite the documented mosquito-repellent properties of *Hyptis suaveolens*, its traditional use relies largely on fresh plant material, which is limited by seasonality and may not provide year-round protection. In addition, evidence on practical, sustained-release delivery systems for plant-derived mosquito repellents remains limited. Candle wax represents a readily available and culturally familiar medium that may facilitate the gradual release of volatile repellent compounds during combustion without requiring direct skin application. This study therefore evaluated the mosquito-repellent efficacy of *Hyptis suaveolens* extracts infused into candle wax as a simple plant-based delivery system. Leaves of *H. suaveolens* were collected, air-dried, ground, and extracted using ethanol and water. The crude extracts were incorporated into melted candle wax at a 5:10 concentration, then solidified and burned in a mosquito-infested room for 30 minutes. Results indicated that rooms with infused candles had significantly fewer mosquitoes than the control room. These findings support the mosquito-repellent properties of Hyptis suaveolens and demonstrate that its incorporation into candle wax is a promising plant-based delivery system for mosquito control, warranting further evaluation under field conditions.

## Introduction

Mosquito-borne diseases represent a significant global health challenge, accounting for nearly 17% of all infectious diseases annually (Brown et al., 2023). Malaria, dengue, Zika virus, and chikungunya continue to place substantial burdens on health systems. Malaria alone was responsible for 229 million cases and over 400,000 deaths in 2019, while dengue accounted for 56 million cases worldwide (Chen et al., 2025; Du et al., 2021; WHO, 2020). The expansion of mosquito populations into new regions due to global trade and travel further complicates control efforts, highlighting the urgent need for effective and sustainable interventions (Abbasi, 2025; Pabst et al., 2025).

Chemical insecticides such as pyrethroids, carbamates, neonicotinoids, and organophosphates have been central to vector control programs (Casida & Durkin, 2013; Davies et al., 2007; WALKER & LYNCH, 2007). Their rapid action against mosquitoes has saved millions of lives, particularly during outbreaks. However, prolonged reliance on these compounds has resulted in significant drawbacks. Insecticide resistance has developed through mechanisms such as metabolic detoxification, target-site mutations, cuticle modifications, and behavioral avoidance (Meier et al., 2022). This resistance, combined with environmental contamination and health risks ranging from neurological damage to endocrine disruption, has led to a pesticide treadmill (Moyes et al., 2017; Namias et al., 2021) that threatens both human and ecological health(Ahmad et al., 2024). Globally, insecticide poisoning affects an estimated 26 million people annually, resulting in approximately 220,000 deaths (Ansari et al., 2014), emphasizing the urgent need for safer alternatives.

In response, research has increasingly focused on plant-derived compounds. Phytochemicals function as repellents, toxins, feeding deterrents, and growth regulators, presenting a promising approach for mosquito control (Koul, 2008; Souto et al., 2021). Citronella, neem, and lemongrass oils have been extensively studied; however, challenges persist regarding standardization, extraction methods, and consistent efficacy (Ali et al., 2023; Choudhury, 2025; Rahaman & Moshwan, 2026). Additionally, most plant-based repellents are formulated as topical applications, which may be costly, inconvenient, or culturally unacceptable in certain communities (Ntonifor et al., 2007).

*Hyptis suaveolens*, locally known in Northern Ghana as “Dumsi nam” (“mosquito rival”), is an aromatic herb traditionally used as a mosquito repellent. Laboratory and field studies have confirmed its mosquitocidal and repellent properties against *Anopheles* and *Aedes* species (Abok et al., 2018; Duniya et al., 2022; Priya et al., 2023) supporting its ethnobotanical use. However, reliance on fresh plant material is constrained by seasonality, as the plant thrives during the rainy season but withers in the dry season (Afreen et al., 2018; Padalia et al., 2015), leaving communities vulnerable when mosquito exposure remains high. Despite its potential, no systematic studies have evaluated *H. suaveolens* extracts in alternative delivery systems capable of providing year-round protection.

This study addresses this gap by evaluating the potential of *Hyptis suaveolens* extract, when infused into candle wax, as a novel mosquito biorepellent. Candles represent a culturally familiar, affordable, and sustained-release delivery system (Purushottam et al., 2024), that can mitigate the seasonal limitations associated with fresh plant use. By comparing repellency across various extraction methods and assessing performance relative to control candles, this research provides new evidence regarding the feasibility of incorporating *H. suaveolens* candles into community-level mosquito control strategies. This approach integrates traditional knowledge with scientific validation and contributes to the development of environmentally safe, accessible, and sustainable alternatives to chemical insecticides.

## Materials and Methods

### Study Design

A mixed-methods approach was used, integrating a community survey with a comparative experimental design to assess the mosquito-repellent efficacy of *Hyptis suaveolens* extract-infused candles. Non-infused candle wax functioned as the negative control, and fresh *H. suaveolens* leaves served as the positive control. A structured questionnaire was administered to 20 residents of Nyankpala Township to document local knowledge and use of *H. suaveolens*. Participants were recruited through convenience sampling and included adults aged 18 to 60 years, representing both genders and a range of occupations, including farming, trading, and student roles.

### Plant Collection and Extraction

Fresh *H. suaveolens* plants were collected from the University for Development Studies (UDS) Nyankpala campus. The plants were cleaned, air-dried for seven days, and pulverized into fine powder. Extractions were conducted at a 1:10 (w/v) ratio by soaking 5g of powdered sample in 50mL of 80% ethanol and distilled water in Falcon tubes. Aqueous extracts were filtered after 24 hours, while ethanolic extracts were filtered after 72 hours. The filtrates evaporated under sunlight, and the crude extracts were weighed and recorded.

### Candle Preparation

Commercial paraffin wax candles were melted using a Delron Single Electric Cooker Hot Plate (Model DE311HA0NVDWJNAFAMZ, Delron Appliances, China), which features a sealed design with thermostat control (AC 220–240 V, 50/60 Hz). Wax was melted at a medium heat setting sufficient to liquefy the wax without burning. For each formulation, 5g of *H. suaveolens* extract was mixed with 10g of melted candle wax, stirred thoroughly, and heated for approximately 15 minutes. The infused wax was poured into 50mL Falcon tubes, cooled to room temperature, and allowed to solidify for 24 hours. The solidified candles were removed from the tubes using a sterile blade. Decoction candles were prepared by frying fresh leaves in melted wax for 15 minutes, followed by cooling, filtering, and drying.

### Experimental Setup

Six identical empty rooms, matched for size, orientation, and corridor location, were prepared. Each room floor was covered with white cloth to make mosquitoes more visible. Windows were sealed to prevent mosquito escape, and doors were left open prior to the experiment to allow natural mosquito entry. Treatments (ethanolic extract candle, aqueous extract candle, decoction candle, non-infused candle, and fresh *H. suaveolens*) were randomly assigned to rooms using a random number generator. Treatment assignments were rotated between replicates to minimize location bias.

Candles were lit simultaneously in all rooms for 30 minutes late at night to ensure comparable ambient conditions. After 15 minutes, the rooms were inspected for dead mosquitoes. The remaining 15 minutes were used to assess repellency, after which candles were extinguished. To standardize mosquito counts, a knockdown was performed using a pyrethroid-based insecticide spray applied uniformly across rooms. Mosquitoes were collected from the cloth floor, counted manually using sterile forceps and placed in labeled containers.

### Addressing Room Bias

To account for potential variation in natural mosquito entry between rooms, treatments were rotated across rooms in successive replicates to minimize location bias. The control room served as the baseline, as no fixed number of mosquitoes was introduced into each room. Repellency was therefore expressed relative to the natural mosquito density present in the control condition. The experiment was repeated three times on different nights, and mosquito counts were averaged across replicates.

### Statistical Analysis

Repellent efficacy was quantified as the Relative Proportion (RP) of mosquitoes recovered in treated rooms compared to non-treatment rooms.

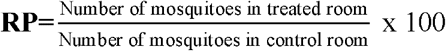

Lower RP values indicated stronger repellency. Data was analyzed using descriptive statistics and comparative proportions to evaluate differences among treatments.

## Results

Four extraction methods of *Hyptis suaveolens* were evaluated for their physical characteristics and mosquito-repellent activity when incorporated into candle wax. Results are presented in both tabular and graphical formats. Table 1 and Figure 1 present the physical properties of the crude extracts, while Table 2 and Figure 3 summarize mosquito recovery after knockdown, expressed as average counts and percentage reductions relative to the control.

**Table 1.** Community knowledge and reported uses of *Hyptis suaveolens* (n = 20).

| ***H. Suaveolens* usage** | **Frequency(n)** | **Percentage (%)** |
| --- | --- | --- |
| Familiar with *H. suaveolens* | 20 | 100 |
| Reported use for mosquito control | 20 | 100 |
| Place fresh plants in rooms | 20 | 100 |
| Burn dried leaves indoors | 13 | 65 |
| Plant around farms | 5 | 25 |

**Table 2.** Physical properties of *Hyptis suaveolens* extracts.

| Extract Type | Texture | Colour |
| --- | --- | --- |
| Ethanolic | sticky | dark green |
| Steam distillation | non-sticky | dark brown |
| Cold water maceration | Non-sticky | dark brown |
| Decoction |  | pale green |

**Fig 1.**
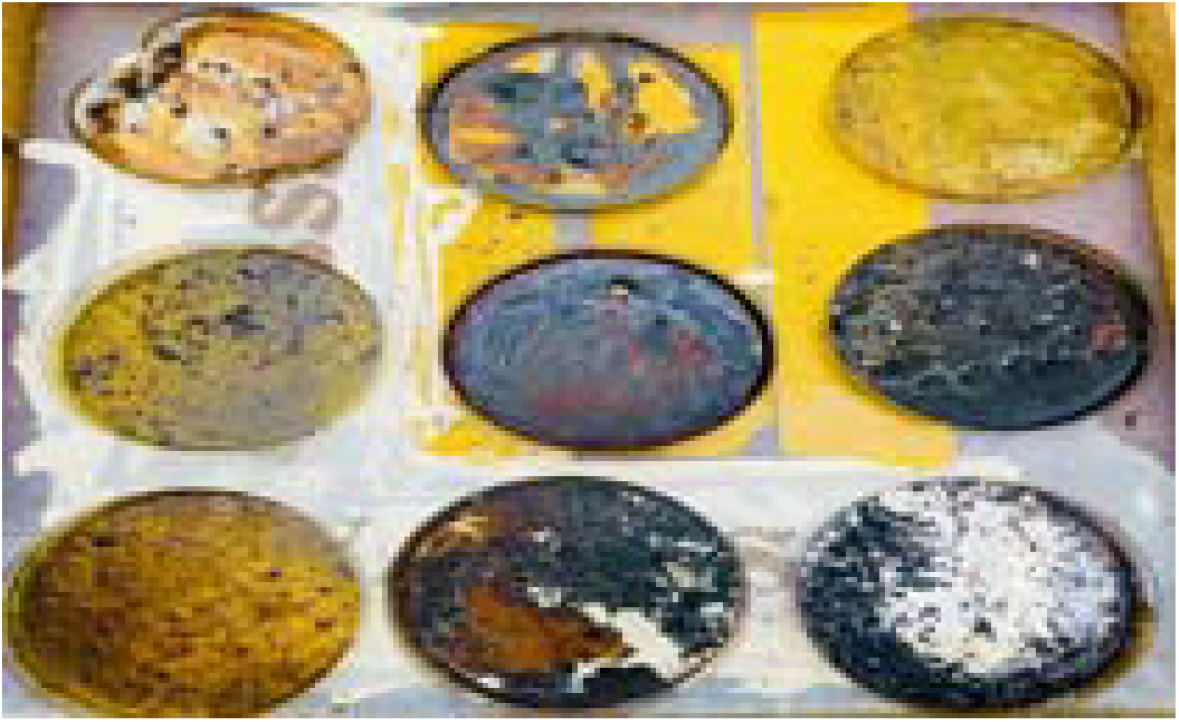
Crude extracts obtained using different extraction methods

**Fig 3.**
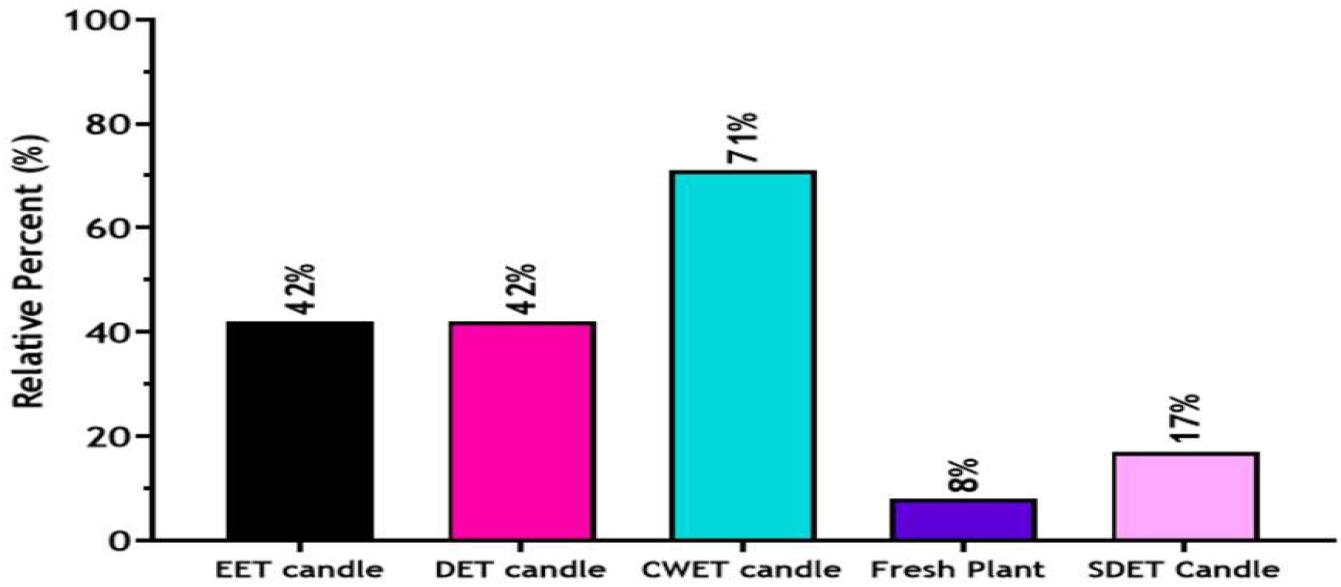
Mosquito repellent activities of extracts-infused candles.

### Community Knowledge and Utilization of *Hyptis suaveolens*

Twenty residents of Nyankpala Township participated in the survey. All respondents were familiar with *Hyptis suaveolens* and acknowledged its mosquito-repellent properties. Every participant also reported using the plant to reduce mosquito nuisance in their homes.

All respondents most frequently placed fresh plants in rooms. Thirteen participants burned dried leaves indoors to repel mosquitoes, and five cultivated the plant around farms to deter insects and other pests. Each participant reported multiple uses.

When asked about their preference for *H. suaveolens*, several respondents cited the high cost of commercial insecticides and repellents. Others reported discomfort, including cold-like symptoms, after using chemical insecticides, prompting them to seek alternatives. Overall, respondents viewed *H. suaveolens* as an accessible, affordable, and effective traditional repellent. Table 1 summarizes respondents’ knowledge of *Hyptis suaveolens* and its reported uses among the 20 participants included in the study.

### Physical Properties of *Hyptis suaveolens* Extracts

The physical characteristics of the *Hyptis suaveolens* extracts differed according to the extraction method. The observed variations in colour and appearance are summarized in Table 2.

Table 2 demonstrates that the extraction method significantly influenced the physical characteristics of the crude extracts. Ethanolic extraction yielded a sticky, dark-green residue, consistent with the solubilization of chlorophylls and polar phytochemicals. Steam distillation yielded a non-sticky, dark-brown extract, indicating a high concentration of volatile oils and terpenoids. Cold-water maceration yielded a non-sticky, dark-brown extract, likely due to the limited solubility of volatile compounds in water. Decoction yielded a pale green extract, suggesting partial pigment degradation and reduced retention of volatile constituents. These findings highlight the importance of the extraction technique in determining the chemical composition and physical properties of *H. suaveolens* extracts.

Figure 1 visually complements Table□1, showing the distinct physical appearances of the four extracts. The image confirms the variations in texture and color described in Table□1, illustrating the sticky dark-green ethanolic extract, the non-sticky dark-brown steam-distilled and cold-water macerated extracts, and the pale-green decoction extract.

### Mosquitoes Recovered After Knockdown Assays

The average number of mosquitoes recovered for each treatment is presented in Table 3. These values were used to calculate relative percentages compared to the control, as illustrated in Figure3.

**Table 3.**
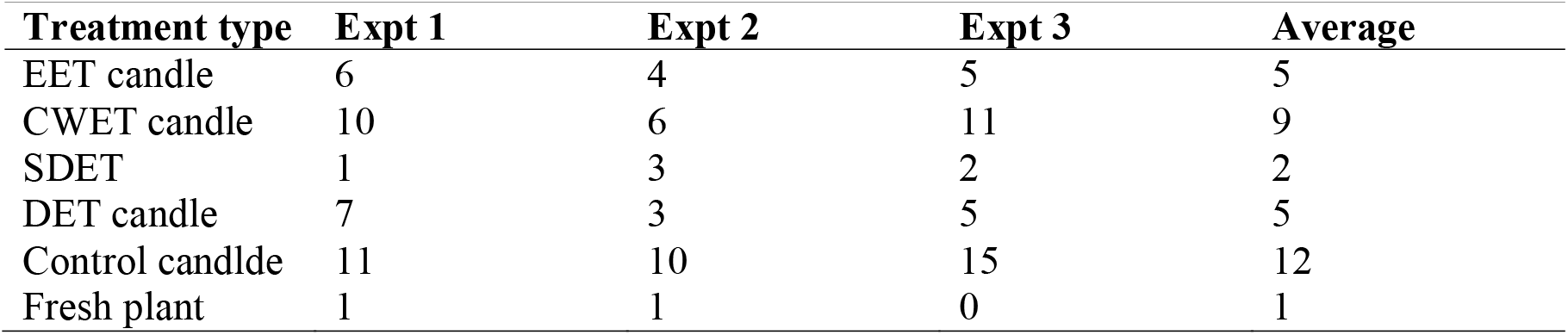
Number of recovered mosquitoes.

| Treatment type | Expt 1 | Expt 2 | Expt 3 | Average |
| --- | --- | --- | --- | --- |
| EET candle | 6 | 4 | 5 | 5 |
| CWET candle | 10 | 6 | 11 | 9 |
| SDET | 1 | 3 | 2 | 2 |
| DET candle | 7 | 3 | 5 | 5 |
| Control candle | 11 | 10 | 15 | 12 |
| Fresh plant | 1 | 1 | 0 | 1 |

EET candle-Ethanol extract candle, SDET Candlle-Steam distillation extract candle, DET candle-Decotion extract candle, CWET candle-Cold water maceration extract candle.

Table 2 indicates that the control candle consistently resulted in the highest mosquito recovery (mean = 12), establishing the baseline density. Fresh plant material was the most effective, with a mean recovery of nearly zero (1). Steam distillation extract candles performed similarly (mean = 2), while ethanolic and decoction candles demonstrated moderate repellency (mean = 5 each). Cold-water maceration was the least effective (mean = 9), approaching control levels.

Note:The x-axis indicates the different extracts-infused candles evaluated, and fresh plant of *Hyptis suaveolens*. The y-axis shows the proportions of mosquitoes recovered after the knocked-down assay, relative to mosquitoes recovered from the test room with a control candle without any extract.

Figure 3 presents the comparative effectiveness of the treatments relative to the control baseline. Fresh plant material was the most effective, reducing recovery to 8% of control counts. Steam distillation extract candles performed similarly, with recovery at 17% of the control. Ethanolic and decoction candles demonstrate moderate repellency, reducing recovery to 42% of the control. Cold-water maceration was the least effective, with recovery at 71% of the control, indicating minimal repellency. These findings indicate that steam distillation and fresh plant use provide the highest and most consistent protection, while cold-water maceration performed poorly.

## Discussion

The use of synthetic insecticides for mosquito control has declined due to limited new products, high costs, environmental and health concerns, and the emergence of resistance (Weng et al., 2024). As a result, eco-friendly strategies such as biological vector control have become increasingly important, with a shift toward plant-based insecticides (Şengül Demirak & Canpolat, 2022). These plant-derived alternatives are characterized by lower toxicity, target specificity, effectiveness at low concentrations, and biodegradability, positioning them as suitable replacements for synthetic compounds. Plants produce phytochemicals that function as repellents, toxins, feeding deterrents, and growth regulators against insects (Vijay Veer & Gopalakrishnan, 2016). The efficacy of these phytochemicals depends on factors such as plant species, plant part, age, solvent polarity, and mosquito species (Shaalan et al., 2005).

In this context, the present study demonstrates that the extraction method significantly influences the mosquito repellent activity of *Hyptis suaveolens* when incorporated into candle wax. Steam distillation and the use of fresh plant material yielded the highest repellency, whereas ethanolic and decoction extracts showed moderate effectiveness, and cold-water maceration was the least effective. These results corroborate previous findings that attribute the mosquito-repellent properties of fresh *H. suaveolens* leaves to volatile compounds (Duniya et al., 2022), and align with studies indicating that solvent polarity and extraction technique impact phytochemical yield and bioactivity (Juhaimi et al., 2019). The superior efficacy of steam distillation highlights the importance of preserving volatile oils, such as terpenoids and essential oils (Mishra et al., 2021), which are widely recognized as mosquito repellents. The reduced effectiveness observed with ethanolic, and decoction methods may be due to the loss or degradation of these volatile compounds.

Utilizing candle wax as a delivery system was both practical and culturally acceptable, providing sustained release of active compounds in indoor environments. Previous studies involving citronella and neem have demonstrated the effectiveness of volatile oils delivered through household products, further supporting the relevance of plant-based repellents as alternatives to synthetic insecticides, which are challenged by resistance, toxicity, and environmental persistence (Weng et al., 2024). Although differences in volatile concentration likely account for the observed variations in efficacy, additional factors such as compound stability, degradation during heating, or interactions with wax may also play a role.

### Limitations

Despite these promising results, several limitations should be acknowledged. The study was conducted in a controlled indoor environment, which may not accurately reflect field conditions. The lack of standardization in the number of mosquitoes per trial could affect reproducibility. Additionally, chemical profiling of the extracts was not performed, limiting the identification of specific active compounds. These constraints reduce the generalizability of the findings and hinder the standardization of formulations.

### Recommendation

Future research should incorporate chemical analyses, such as gas chromatography–mass spectrometry (GC–MS), to identify and quantify active constituents. Long-term field trials are necessary to evaluate real-world effectiveness, and studies on community acceptance are needed to assess potential adoption. Comparative investigations across mosquito species and environmental conditions would strengthen the evidence base, while optimization of extraction methods and candle formulations could improve efficacy and scalability.

## Conclusion

The results indicate that *Hyptis suaveolens* extracts incorporated into candle wax effectively repel mosquitoes, with efficacy dependent on the extraction method. Candles infused with steam-distilled extracts achieved the highest repellent activity, reducing mosquito counts by up to 83% compared to control rooms. On average, rooms with extract-infused candles exhibited a 58% reduction in mosquito counts. Fresh plant material also demonstrated strong repellency, highlighting the significance of volatile compounds. These findings align with previous reports that phytochemical yield and bioactivity are influenced by extraction technique and solvent polarity. Furthermore, the results highlight the practical utility of candle wax as a culturally acceptable delivery system for the sustained release of active compounds.

Overall, these results emphasize the potential of plant-based mosquito repellents as safe, affordable, and environmentally sustainable alternatives to synthetic insecticides, which are increasingly constrained by resistance, toxicity, and ecological persistence. *Hyptis suaveolens* extracts, especially those obtained through steam distillation, offer a promising strategy for household mosquito control in resource-limited environments.

